# TOMVi: A Tool for Overviewing Metadata Visualization

**DOI:** 10.1101/506220

**Authors:** Daisuke Kyoui, Hiroyuki Yamamoto, Natsumi Mikami, Ryota Ishii, Hiroki Tanaka, Naruki Fukuda, Taketo Kawarai, Hirokazu Ogihara

**Affiliations:** Laboratory of Food Hygiene, Department of Food Bioscience and Biotechnology, College of Bioresource Sciences, Nihon University.

## Abstract

DNA sequencing technology has been improved drastically in the recent decades, which allows comprehensively analyze microbial community and gene expression. Simultaneously with the improvements, many tools and software for much amount of sequence data have been released. These analysis tools/software can produce statistic information. However, to interpret and to find the relationships and/or associations between the information and the experimental condition may be a complex task, especially for non-bioinformaticians. In this article, an open source software TOMVi (Tool for Overviewing Metadata Visualization) which visualized similarity and relationships between the samples corresponding to the experimental conditions was introduced. TOMVi allows researcher to interactively manipulate the composition data in each sample toward to each experimental condition such as location, time-series and others with graphical user interface (GUI), and to visualize the existence and behavior of each OTU. The figure illustrated by this tool can provide an intuitive suggestion but not always correct as statistically, and will be helpful to decide further analyzing tactics.

The TOMVi is available at http://github.com/Daisuke-Kyoui/tomvi.

## Background

In this decade, next generation sequencing (NGS) became common, and allows researchers to obtain extraordinary much DNA sequence (Levy et al. 2016). This technology has been applied to various approaches to elucidate organism in genomics, transcriptomics and omics. Furthermore, NGS technologies has been improving forward to more much, long and accurate output.

Metagenome analysis is one of the most matured approach among the approaches using NGS. Because of this approach requires much amount of reads, NGS which can produce short (approximately 50∼300 bp) and many reads (over mega-reads) was very suitable for the purpose. Metagenome analysis mainly targets the microbes in the environment, and reveals the community and metabolisms in the sample (Tyson et al. 2004). Especially, metagenome analysis targeting 16S rRNA amplicon which links to taxonomic position became common approach to reveal the microbial community in human, food and other natural environments (Caporaso et al. 2011).

To interpret the metagenome sequencing results, many tools and software are released. Most of them were developed by bioinformaticians with discussions on the web (Caporaso et al. 2010). These tools and software analyze much amount of sequences statistically, and provide to researchers as intensive information. Although the tools and software provide intensive information, the information is still large and complex. Moreover, for the researcher who is not familiar with bioinformatics skill, to interpret the sequencing results by comparing numerical data may be a big obstacle.

TOMVi (Tool for Overviewing Metadata Visualization) was developed for the researchers who is not a specialist of bioinformatics. On this tool, researcher can interactively manipulate plots which show composition of OTUs in each samples toward to corresponding experimental condition by graphical user interface (GUI). Then, this tool visualizes behavior of OTU rates between the plots as a line width change which connecting experimental conditions. Finally, TOMVi provides overviewing of increase or decrease of OTUs between experimental conditions. The figure which obtained by using TOMVi is not show the relationships statistically between the samples like PCoA, rather describes relationships between the OTU composition and experimental condition. Briefly, the figure can be provide an intuitive suggestion to researcher who is needed to decide which OTUs should be paid attention or how proceed analyze more.

## TOMVi

TOMVi is a thoroughly tested and open-source tool which implemented with HTML5 and Javascript, with plugins (jCanvas: https://github.com/caleb531/jcanvas; jQuery: https://github.com/jquery/jquery). TOMVi requires two input files. One is the “background file” which is a graphical file formatted in jpeg or png. This file shows experimental condition such as location, time-scale, temperature and anything, and can be designed by user’s purpose. The other one is “datafile” which is a tab deliminated table like a heatmap. In the table, the header lists samples, the left end column lists OTUs and cells show the rate of OTU in each sample. Optionally, “links list” file can be loaded. This file lists links between the samples which are required hidden or visible, for the purpose of such as time-series visualization. After loading files, samples are visualized as a plot of circle graph which shows the rates of OTUs in the sample, and are ordered on the background. Then, user can drag each plot to desirable position on the background figure, then re-draw links between the plots. The links are colored corresponding to each OTU. And its width at connecting to plot indicates the rate of OTU in the plot. Finally, on the background, relationships between samples are visualized. Moreover, if user tracing specific OTU, the others can be hidden. And also, user can compare the rate of each OTUs between samples. All of the manipulation of TOMVi can be controlled by GUI, which may be friendly for the user who is not familiar to command-line user interface (CUI).

## Case study

As an example, the dynamics of microflora during spoilage of fish was visualized (Figure 1). In this case, jack mackerel (*Trachurus japonicus*) was stored until 48 h and its microflora at skin, muscle and viscera were analyzed respectively by metagenome analysis targeting 16S rRNA. Figure 1 was generated by TOMVi, and its background illustrated a cross section of jack mackerels. Figure 1A shows the behavior of all OTUs in which of minimum rate was more than 0.01%. This overview described the microflora transition during spoilage. At 0h which means before storage, the OTUs colored from blue to purplish red were dominant OTUs. Contrastingly, from 24h storage, OTUs colored from green to light green became dominant. Also, from 24h to 48h, oligopolization could be noticed. Figure 1B was made from same data with Figure 1A. But, all OTUs except of “denovo8364” which is the most dominant OTU at 48h-muscle were hide. This figure showed the behavior of the denovo8364 which was becoming dominant from 24h storage and was growing at muscle only. Summary, in this case, TOMVi describes the overview of the dynamics of microflora intuitively and suggested which OTUs might have a major role in jack mackerel spoilage. These observation could be helpful for further statistical analysis or examination such as isolating the species.

**Figure 1.**
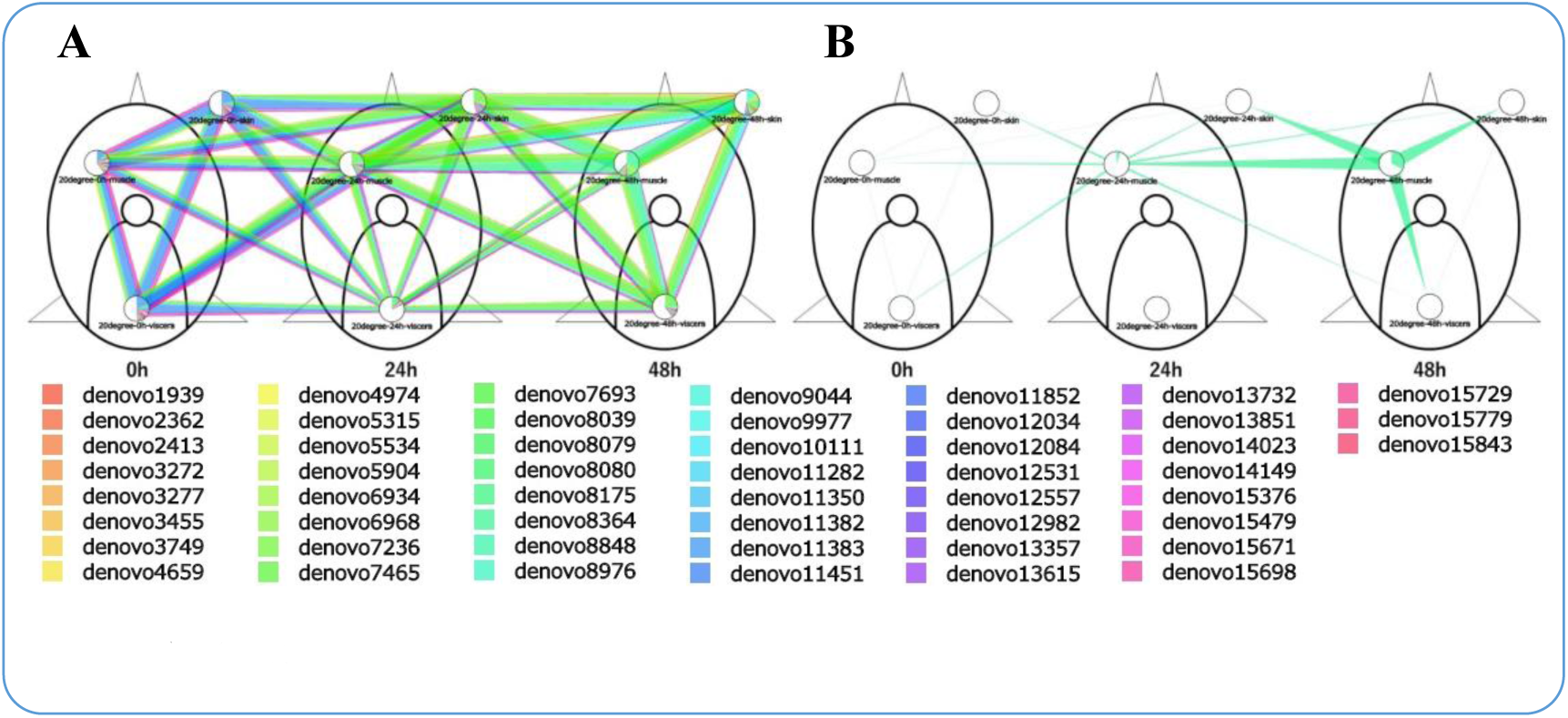
The microbiota dynamics during jack mackerel spoilage generated by TOMVi. A) The OTUs in which of rate was more than 0.01% were visualized. B) Only “denovo8364” which is the most dominant OTU in 48h-muscle was visualized.

## Availability of source code and requirements

Project name: TOMVi

Project home page: http://github.com/Daisuke-Kyoui/tomvi

Programming language: HTML5, JavaScript and CSS

Browsers: Chrome, Safari and FireFox were tested.

Plugins: jQuery-3.3.1 and jCanvas are used.

License: MIT license

No restrictions of use for academic or non-academic purposes.

